# Loss of cohesin subunit Stag1 in zebrafish limits cell cycle progression and is compensated by altered BMP signalling and metabolic pathways

**DOI:** 10.64898/2026.08.20.745648

**Authors:** Dylan M. Lynch, Anastasia A. Labudina, Sarada Ketharnathan, Rachel Coldicott, Christoph Goebl, Julia A. Horsfield, Michael Meier

## Abstract

Cohesin is a large multisubunit protein complex that plays essential roles in cell proliferation, genome organisation, and gene regulation in metazoans. Germline mutations in cohesin subunits or regulators cause a group of human developmental disorders collectively known as cohesinopathies. Increasing evidence indicates that individual cohesin subunits can confer distinct molecular functions to the complex; for example, STAG1 and STAG2 have both overlapping and non-overlapping roles in genome organisation. The zebrafish tailbud provides an excellent developmental model for investigating the coordination of cell proliferation and differentiation; processes in which cohesin has crucial functions. We previously demonstrated that loss of Stag2 disrupts Wnt signalling and mesoderm patterning in the zebrafish tailbud. Here, we show that, unlike mammals, zebrafish can tolerate complete loss of Stag1 from embryogenesis through to adulthood. In contrast to Stag2 deficiency, loss of Stag1 impairs cell cycle progression, activates p53 signalling, and induces a metabolic shift towards catabolism. BMP signalling is reduced in Stag1-deficient embryos and is accompanied by expansion of BMP antagonist *chordin* expression. Stag1 loss also alters chromatin accessibility at the *chordin* locus and affects accessibility at chromatin domain boundaries. We propose that modulation of growth and signalling pathways compensates for the absence of Stag1, allowing embryonic development to proceed correctly. Together, these findings reveal distinct contributions of Stag1 and Stag2 to cell-cycle regulation, chromatin architecture, and developmental signalling during vertebrate embryogenesis.

## Introduction

Cohesin is a highly conserved multiprotein complex essential for chromosome segregation, DNA repair and gene regulation reviewed in (Rittenhouse & Dowen, 2024). It consists of a core ring (SMC1, SMC3 and RAD21) and two mutually exclusive associated HEAT repeat-containing protein subunits (either STAG1 or STAG2) (Sumara et al., 2000). In eukaryotes, the ATP-dependent DNA loop extrusion activity of the cohesin complex plays a central role in the three-dimensional (3D) folding of chromosomes in interphase. Additional factors regulate extrusion rates in a dynamic fashion by either constantly loading (heterodimer of NIPBL-MAU2) (Petela et al., 2018), or releasing (PDS5A/B and WAPL) cohesin from chromatin (Kueng et al., 2006). Cohesin is retained and stabilised when it encounters the N-terminal part of CTCF (Pugacheva et al., 2020) forming loop anchors (Rao et al., 2014). The result is a CTCF-anchored partitioning of the genome into topologically associated domains (TADs), which have a ∼2-3 fold higher interaction frequency compared to regions outside the TAD measured by Hi-C (Dixon et al., 2012; Lieberman-Aiden et al., 2009). This higher order chromatin structure is thought to have a gene-regulatory function by facilitating genomic interactions between enhancers and promoters inside TADs while at the same time restricting interactions across TAD boundaries (Chakraborty et al., 2025; Lupianez et al., 2015). While the two STAG-cohesin complexes have largely overlapping roles in sister chromatid cohesion, they exhibit divergent functions in interphase genome organisation and gene regulation (Casa et al., 2020; Chang et al., 2026; Cuadrado et al., 2019; De Koninck et al., 2020; Ketharnathan et al., 2020; Kojic et al., 2018; Viny et al., 2019; Wang et al., 2026). STAG2-cohesin has been extensively studied because of the high frequency of *STAG2* mutations in human cancers reviewed in (Scott et al., 2025). In contrast, the developmental roles of STAG1-cohesin remain poorly defined, albeit variants identified in human studies have shown to cause unspecific syndromic intellectual disability (Lehalle et al., 2017; Zhang et al., 2025).

STAG1- and STAG2-cohesin display distinct genomic distributions and architectural functions. STAG1-cohesin is more resistant to WAPL-mediated release enabling the formation of longer and more stable loops often found at CTCF-bound sites (Kojic et al., 2018; Wutz et al., 2020). STAG2-cohesin is more abundant in general, about 75% of the cohesin pool, at non-CTCF sites and contributes to cell-type-specific shorter (<100 kbp) enhancer-promoter contacts and polycomb domain organisation (Cuadrado et al., 2019).

Despite these molecular differences, single *Stag* mutants are viable in cell culture. In human cancer cell lines, individual STAG1 or STAG2 knockout is tolerated, whereas simultaneous loss of both subunits phenocopies core cohesin depletion and is lethal (van der Lelij et al., 2017). In mice, constitutive *Stag1* or *Stag2* knockouts are embryonic lethal (Cuadrado et al., 2015; De Koninck et al., 2020), but the phenotypes differ from the catastrophic mitotic failure caused by loss of core cohesin subunits and instead reflect developmental defects. Conditional *Stag2* deletion in adult mice is viable but impairs haematopoiesis, further indicating that STAG proteins have essential functions outside of mitosis (Viny et al., 2019). These observations suggest that each Stag subunit performs non-redundant roles in interphase that are important for development and tissue homeostasis.

The zebrafish model offers a powerful system to dissect these subunit-specific functions. Single *stag1b* and *stag2b* mutants are viable and fertile, whereas simultaneous loss of both paralogues phenocopies the mitotic catastrophe observed in *rad21* mutants (Horsfield et al., 2007; Labudina et al., 2024). This genetic separation enables investigation of Stag1- and Stag2-specific contributions to embryonic development without the confounding effects of mitotic failure. Here we use the rapidly proliferating and differentiating population of embryonic cells in the zebrafish tailbud to show that Stag1 loss, unlike loss of Stag2, impairs cell cycle progression, disrupts BMP signalling through expansion of the *chordin* expression domain, and weakens chromatin accessibility specifically at TAD boundaries.

## Results

### Stag1b deficiency has shared and unique consequences for gene expression compared with loss of Stag2b in developing zebrafish tailbuds

We previously created zebrafish mutants for genes that encode the Stag paralogues, Stag1a, Stag1b, and Stag2b (Ketharnathan et al., 2020) and Rad21 (Horsfield et al, 2007). *rad21^-/-^* mutants are embryonic lethal and die by mitotic catastrophe, with cells that cannot complete mitosis. Double homozygous mutation in *stag1a;stag1b* is also lethal and phenocopies *rad21^-/-^* homozygotes (Labudina et al., 2024), indicating that the *stag1a* and *stag2a* paralogues cannot compensate for the loss of *stag1b* and *stag2b*. The viability of single *stag* mutants affords the opportunity to explore the individual functions of Stag subunits in development. Previously we showed that Stag2b is important for mesoderm cell fate in the zebrafish tailbud, and that loss of Stag2b downregulates Wnt signalling. However, it is unclear how the Stag1 paralogues influence development in zebrafish, and if this differs from Stag2b.

We performed bulk RNA sequencing (RNA-seq) on tailbuds from genotypes *stag1b^-/-^*, *stag2b^-/-^* and *rad21^-/-^* at 16 hours post-fertilisation (hpf) as previously described (Labudina et al., 2024) (Fig. 1A). We found 1,972 significantly differentially expressed genes (DEGs) in *stag1b^-/-^* (988 downregulated, 984 upregulated) and 948 DEGs in *stag2b^-/-^* (560 downregulated, 388 upregulated) respectively (*Padj.* < 0.05) (Fig. 1B, Data S1, S2). 293 DEGs overlapped in both genotypes (Fisher Exact Test, *P* < 0.00001, OR 5.19).

**Figure 1.**
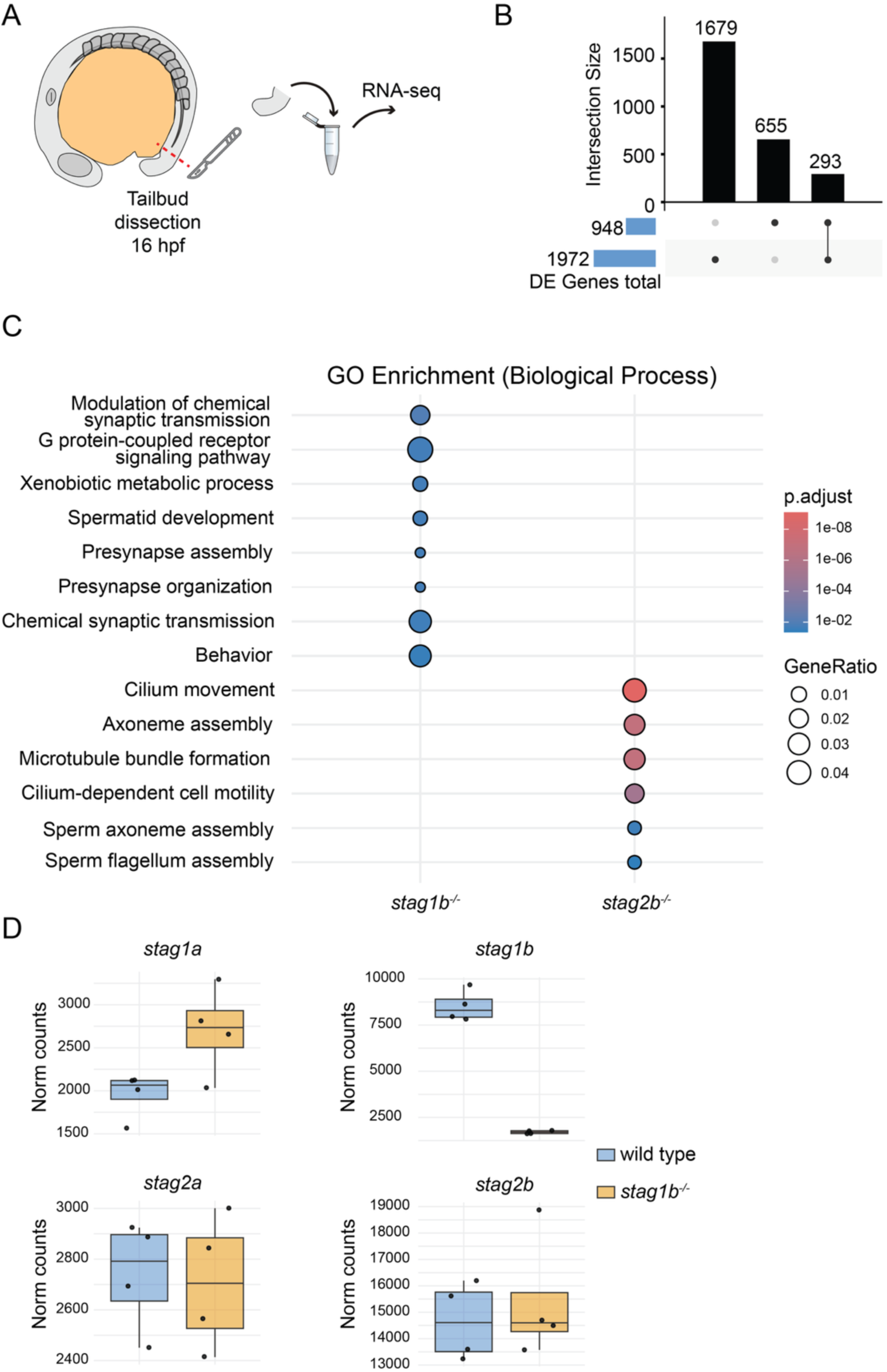
RNA-seq analysis of *stag1b^-/-^* and *stag2b^-/-^* mutants reveals shared and distinct effects on gene expression. (**A**) 40 tailbuds for each of four biological replicates of wild type, *stag1b^-/-^* and *stag2b^-/-^* mutants were dissected and processed for RNA-seq. (**B**) Upset plot showing common and distinct differentially expressed genes (DEGs) in *stag1b^-/-^* and *stag2b^-/-^*mutant tailbuds. (**C**) Gene ontology enrichment analysis of DEGs in *stag1b^-/-^* and *stag2b^-/-^* mutants. (**D**) Normalised read counts for *stag* paralogues from *stag1b^-/-^* and wild type tailbuds.

We conducted gene ontology (GO) enrichment analysis for biological processes of significant DEGs in both genotypes (Fig. 1C, Data S3). This analysis showed that *stag1b^-/-^* tailbuds were significantly enriched for terms related to chemical synaptic transmission, regulation of trans-synaptic signalling, G-protein-coupled receptor signalling and xenobiotic metabolic process. In contrast, the *stag2b^-/-^* tailbuds were strongly enriched for processes associated with cilium movement, axoneme assembly, microtubule bundle formation and cilium-dependent cell motility. RNA-seq analysis also showed an increase in *stag1a* expression, potentially compensating for the loss of *stag1b* (Fig. 1D).

Complete loss of Stag1 is tolerated in zebrafish development but affects cell cycle progression. To determine if Stag1a compensates for Stag1b in *stag1b^-/-^* mutants, we crossed *stag1a^-/-^* fish (Ketharnathan et al., 2020) with *stag1b^-/-^* to create *stag1a;stag1b* double homozygous mutants. Surprisingly, these fish are viable and show no gross morphological defects, although their development is slightly delayed (Fig. 2A). Therefore we propose that in zebrafish, other Stag subunits can largely compensate for loss of Stag1 in development.

**Figure 2.**
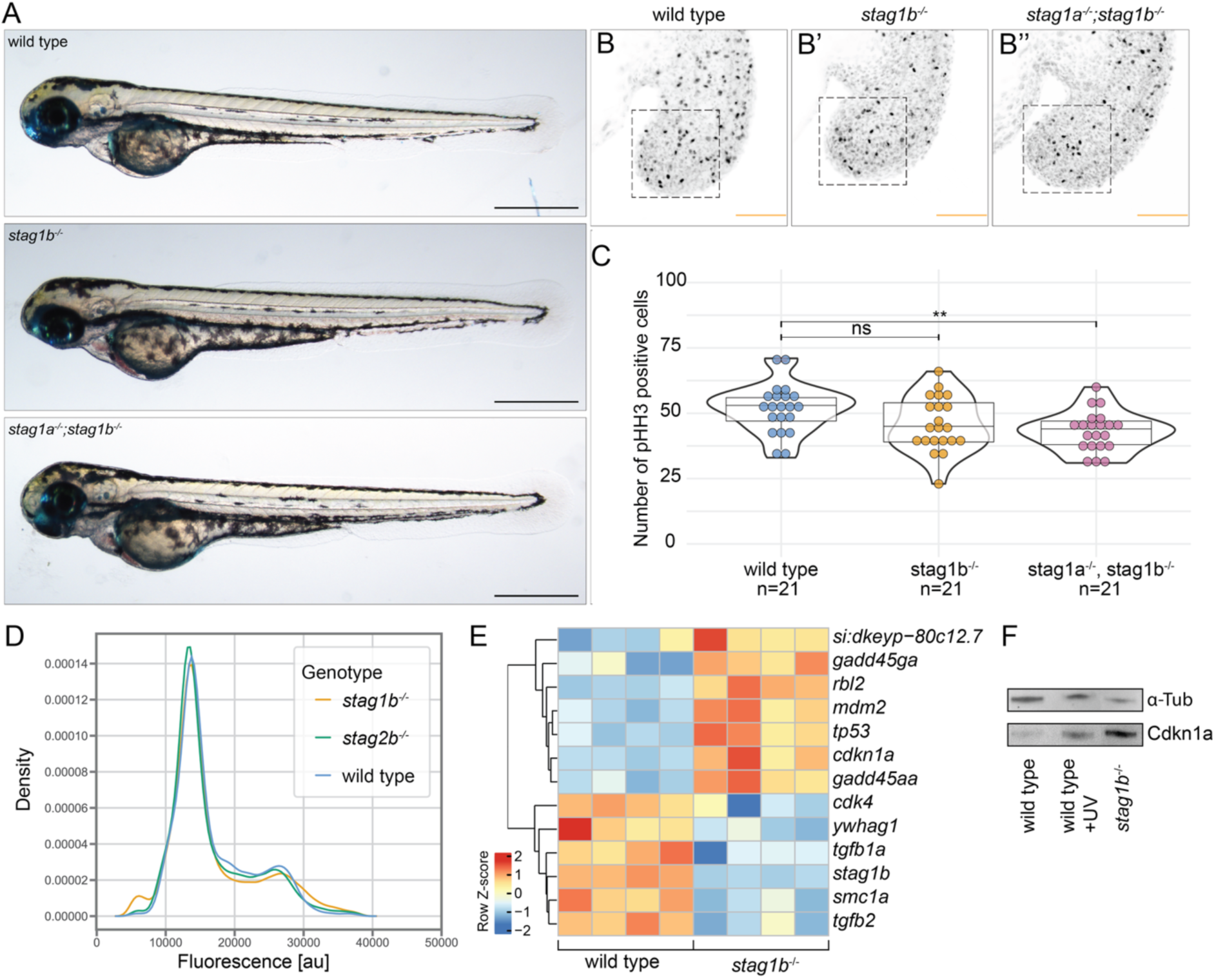
Stag1 loss is tolerated in zebrafish development but leads to developmental delay and upregulation of *tp53*. (**A**) Lateral view of zebrafish larvae at 3 days post fertilization (dpf) for genotypes wild type (top), *stag1b^-/-^* (middle) and *stag1a^-/-^;stag1b^-/-^* (bottom). Scale bars are 500 µm. (B) pHH3+ cells in 16 hpf wild type (**B**), *stag1b^-/-^* (**B’**) and *stag1a^-/-^;stag1b^-/-^* (**B’’**) tailbuds. (**C**) Violin and box plots of pHH3 positive cells were counted in the area corresponding to the dashed box in (**B**) (n = 21 tailbuds per genotype). Statistical significance was determined using a t-test: ** p ≤ 0.01, ns = not significant. (**D**) Cytometry cell cycle profiles of wild type, *stag1b^-/-^*and *stag2b^-/-^* tailbud cells. (**E**) Heatmap of DEGs involved in cell cycle regulation in *stag1b^-/-^* mutants compared to wild type. (**F**) Western blot detecting p21 (encoded by *cdkn1a*) and alpha-tubulin (α-Tub) in wild type, UV treated and *stag1b^-/-^* mutant whole zebrafish embryos (n=30 embryos per condition and genotype).

To investigate the delayed growth upon loss of Stag1, we performed whole mount immunohistochemistry detecting phosphorylated histone H3 (pHH3+) marking cells in the G2/M phase of the cell cycle in tailbuds at 16 hpf. There were significantly fewer pHH3+ cells in *stag1a;stag1b* double mutants and there were also fewer cells in *stag1b^-/-^* (although not statistically significant) (Fig. 2B-B” and C)

We performed flow cytometry on cells isolated from *stag1b^-/-^*tailbuds but found no major differences in cell cycle profiles. (Fig. 2D). Nevertheless, genes required for DNA replication, G1/S and G2/M progression, and mitosis are significantly downregulated in *stag1b^-/-^* tailbuds (Fig. 2E). In parallel, key stress-response and cell cycle arrest genes, most notably *tp53* and p21 encoded by *cdkn1a* are strongly upregulated (Fig. 2E). To test whether the transcriptional induction of *cdkn1a* resulted in elevated protein levels, we performed Western blot analysis on 16 hpf whole embryo lysates. p21 protein was markedly increased in *stag1b^-/-^* embryos relative to wild-type controls (Fig. 2F, Supplementary Fig. 1). As a positive control for p53-pathway activation, UV irradiation of wild-type embryos also induced p21. RNA-seq analysis also revealed downregulation of the TGF-β ligands *tgfb1a* and *tgfb2* (Fig. 2E). TGF-β1 and TGF-β2 generally function as negative regulators of cell cycle progression in many cell types, inducing G1 arrest through upregulation of cyclin-dependent kinase (CDK) inhibitors such as p21. Their downregulation in *stag1b^-/-^*tailbuds may therefore represent a compensatory mechanism aimed at restoring proliferation to partially counteract a strong p21-mediated cell cycle arrest.

### BMP signalling is attenuated in a Stag1 dose-dependent manner

Although BMP signalling itself was not significantly enriched in the Gene Ontology analysis, several core components of the pathway, including the BMP antagonist *chordin* (*chrd*) and *follistatin-like 1a* (*fstl1a*) were upregulated in *stag1b^-/-^* tailbuds. Given the central role of BMP signalling in tailbud patterning and its known interplay with cell-cycle control, we next examined this pathway in more detail. RNA-seq analysis showed downregulation of several BMP ligands (*bmp3, bmp4, bmp6*) and the downstream effector *id3* in *stag1b^-/-^* (Fig. 3A). We independently validated some of these genes by RT-qPCR in *stag1b^-/-^*, *stag1a;stag1b* double mutant as well as *stag2b^-/-^* mutants at 16 hpf. We detected a significant dose-dependent upregulation of *chrd* and *fstl1a* in *stag1* deficient embryos (Fig. 3B, C). Importantly, the upregulation of BMP antagonists was specific to *stag1* mutants and did not occur in *stag2b^-/-^* embryos (Fig. 3B, C). BMP ligands such as *bmp4* were significantly reduced in *stag1b^-/-^* and trending lower in double mutants but were not affected in *stag2b^-/-^* (Fig. 3D). *id3* was not significantly changed but also trended lower in *stag1* mutants (Fig. 3E).

**Figure 3.**
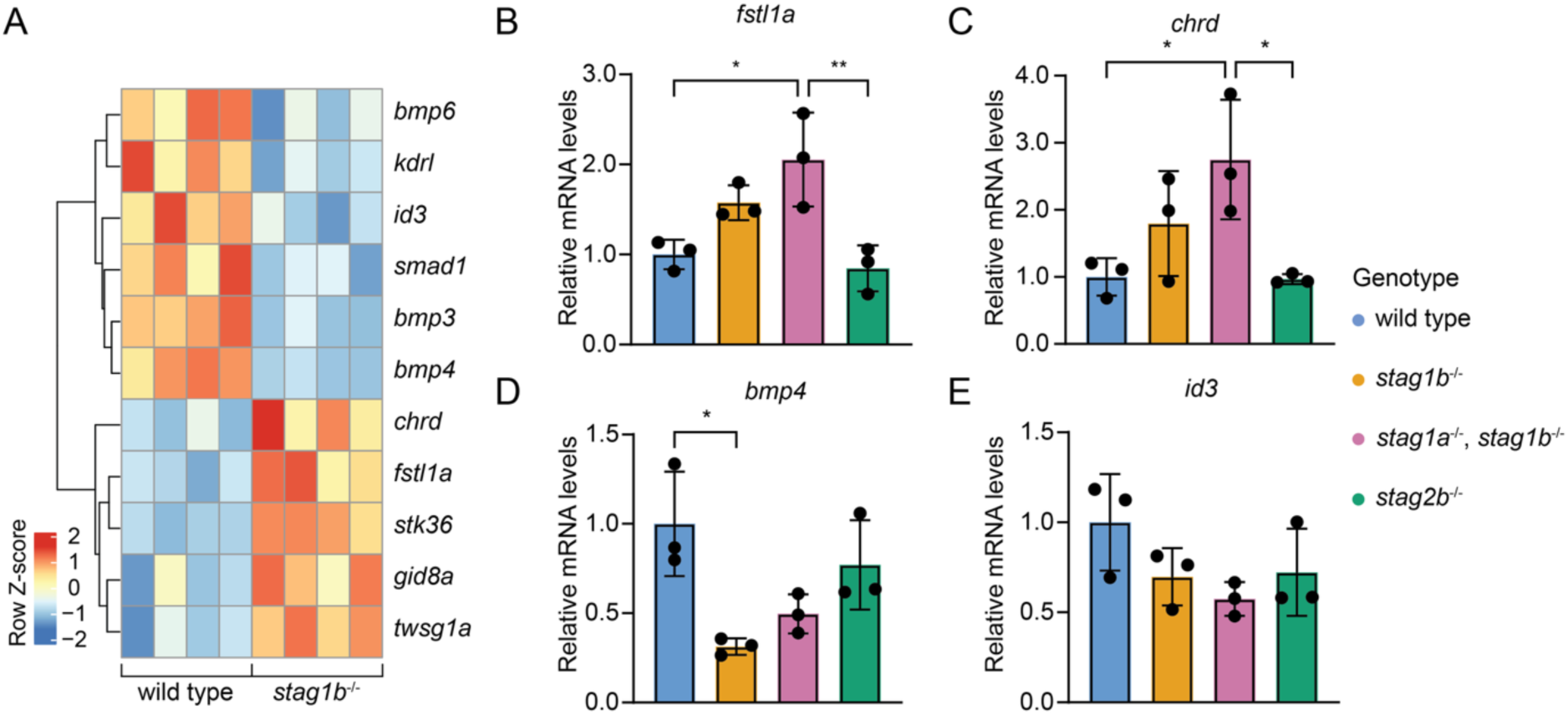
Genes in the BMP signalling pathway are disrupted in *stag1b^-/-^* and *stag1a^-/-^*; *stag1b^-/-^*mutant tailbuds. (**A**) Heatmap of DEGs involved in the BMP pathway in *stag1b^-/-^* mutants compared to wild type showing upregulation of BMP antagonists *chrd* and *fstl1a* as well as downregulation of *bmp3 bmp4* and *id3*. (**B**-**E**). Quantitative RT-PCR of *fstl1a, chrd, bmp4* and *id3* comparing wild-type, *stag1b^-/-^, stag1a^-/-^/stag1b^-/-^* and *stag2b^-/-^* tailbuds. Statistical significance was determined using one-way ANOVA test: * = p ≤ 0.05, ** = p ≤ 0.01.

To determine whether the quantitative changes in BMP pathway gene expression were reflected in altered spatial expression patterns, we performed whole mount *in situ* hybridisation detecting mRNA for the BMP inhibitor *chrd* and the ligand *bmp4* (Fig. 4). In wild-type embryos, *chrd* is expressed just anterior of the mesodermal progenitor zone marked by *bmp4* (Fig. 4 A,B). In *stag1b*^-/-^ mutants (47/50) this domain is expanded into neighbouring cells to the anterior (Fig. 4A’). *stag1a;stag1b* double mutants (48/49) show an even more pronounced expansion of *chrd* (Fig. 4A’’). Conversely *bmp4* is markedly reduced in *stag1b^-/-^* (Fig. 4B’) and double mutants (Fig. 4B’’), although only around half of the analysed embryos showed this phenotype. *stag2b^-/-^* showed no distinct change in either *chrd* or *bmp4* spatial expression or intensity (Fig. 4A’’’ and B’’’). Expansion of *chrd* was not due to developmental delay in *stag1* mutant embryos (Supplementary Fig. 2). Together these results indicate that *stag1a* is functioning additively in the same pathway as *stag1b* to maintain normal expression of BMP antagonists.

**Figure 4.**
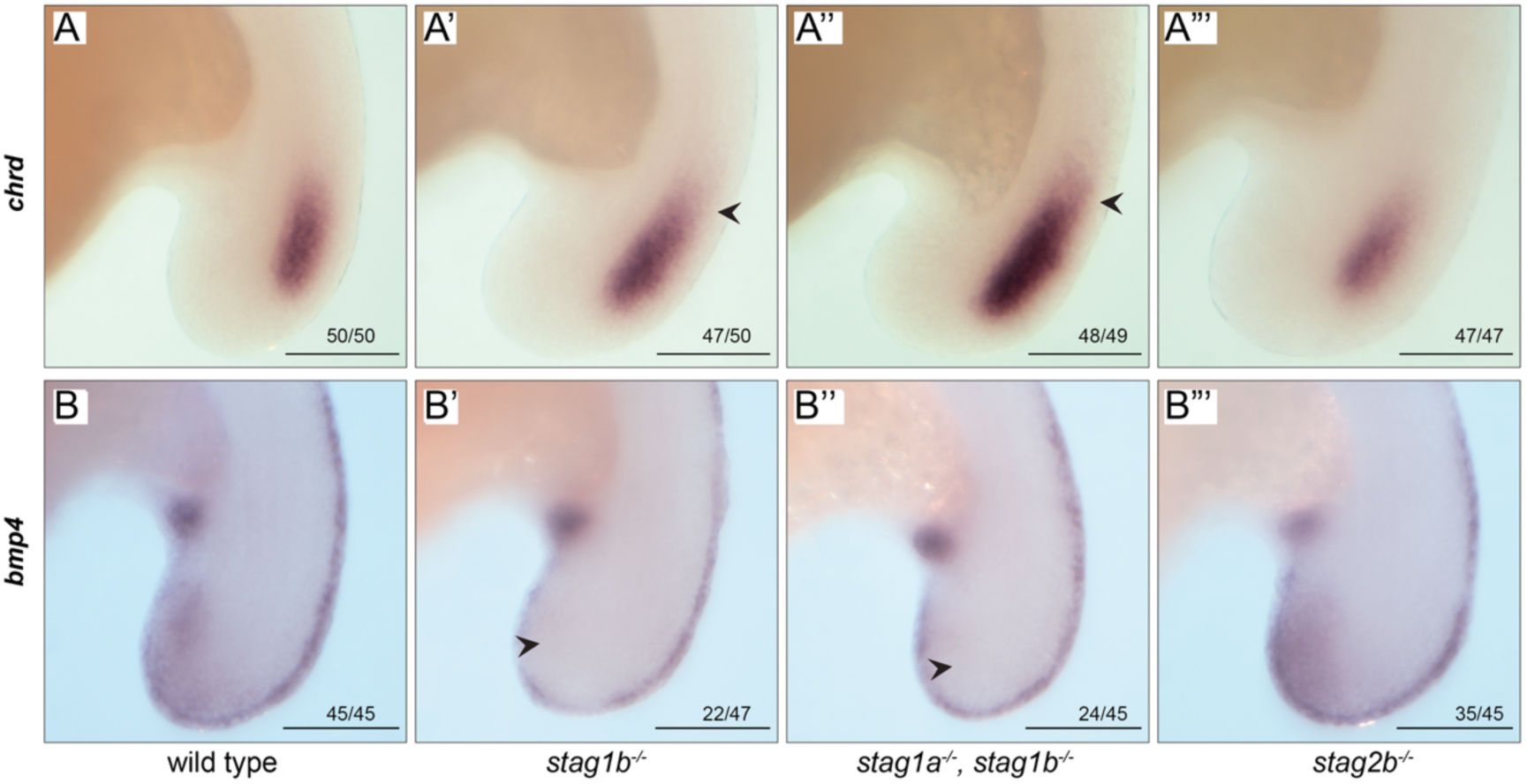
Spatial expression of *chrd, and bmp4* is altered in a stag1 dose dependent manner in 16 hpf zebrafish embryos. Whole mount *in situ* hybridisation showing *chrd* (**A-A”’**) and *bmp4* (**B-B”’**) expression in wild-type, *stag1b^-/-^, stag1a^-/-^;stag1b^-/-^* and *stag2b^-/-^* embryos. *chrd* is expanded into neighbouring cells in both *stag1b^-/-^* and *stag1a^-/-^;stag1b^-/-^* mutants (arrowheads), with a stronger effect in the double mutant. *bmp4* is reduced in the tailbud of *stag1* mutants (arrowheads), again in a dose-dependent manner. *stag2b^-/-^* embryos show no obvious change in *chrd* or *bmp4* expression in the tailbud. Each panel represents a representative photograph with numbers of embryos displaying the representative phenotype at the bottom right (n > 20 embryos per genotype). Scale bar 250 μm.

### *Stag1* loss leads to a dose-dependent increase in chromatin accessibility downstream of the *chordin* locus

To test if the global transcriptional changes and in particular at BMP regulators were caused by changes in the chromatin structure we performed an Assay for Transposase-Accessible Chromatin using sequencing (ATAC-seq) experiment on 16 hpf isolated tailbuds from wild type, *stag1b^-/-^*and *stag1a;stag1b* double mutant embryos. At the *chrd* locus, ATAC-seq revealed increased chromatin accessibility in *stag1* mutants both near the gene itself and at a prominent intergenic region located approximately 90 kb downstream of the transcription start site (Fig. 5A, Supplementary Fig. 3). Smaller increases in accessibility were observed around the *chrd* promoter and nearby intergenic regions in *stag1* mutants. However, a much stronger *stag1* dose-dependent gain was detected at the distal intergenic region between *ilkap* and *clcn2c* (Fig. 5A,B). The region lies between *clcn2c* (a voltage-gated chloride channel) and *ilkap* (a serine-threonine phosphatase inhibiting senescence and cell death). *clcn2c* was significantly upregulated in *stag1b^-/-^* mutant tailbuds (Data S1). *Ilkap* was also trending higher, albeit not significant, suggesting de-repression of a cluster of genes in the neighbourhood of *chrd*. This distal peak was most pronounced in *stag1a;stag1b* double mutants, consistent with the dose-dependent expansion of the *chrd* expression domain observed by in situ hybridisation (Fig. 4).

**Figure 5:**
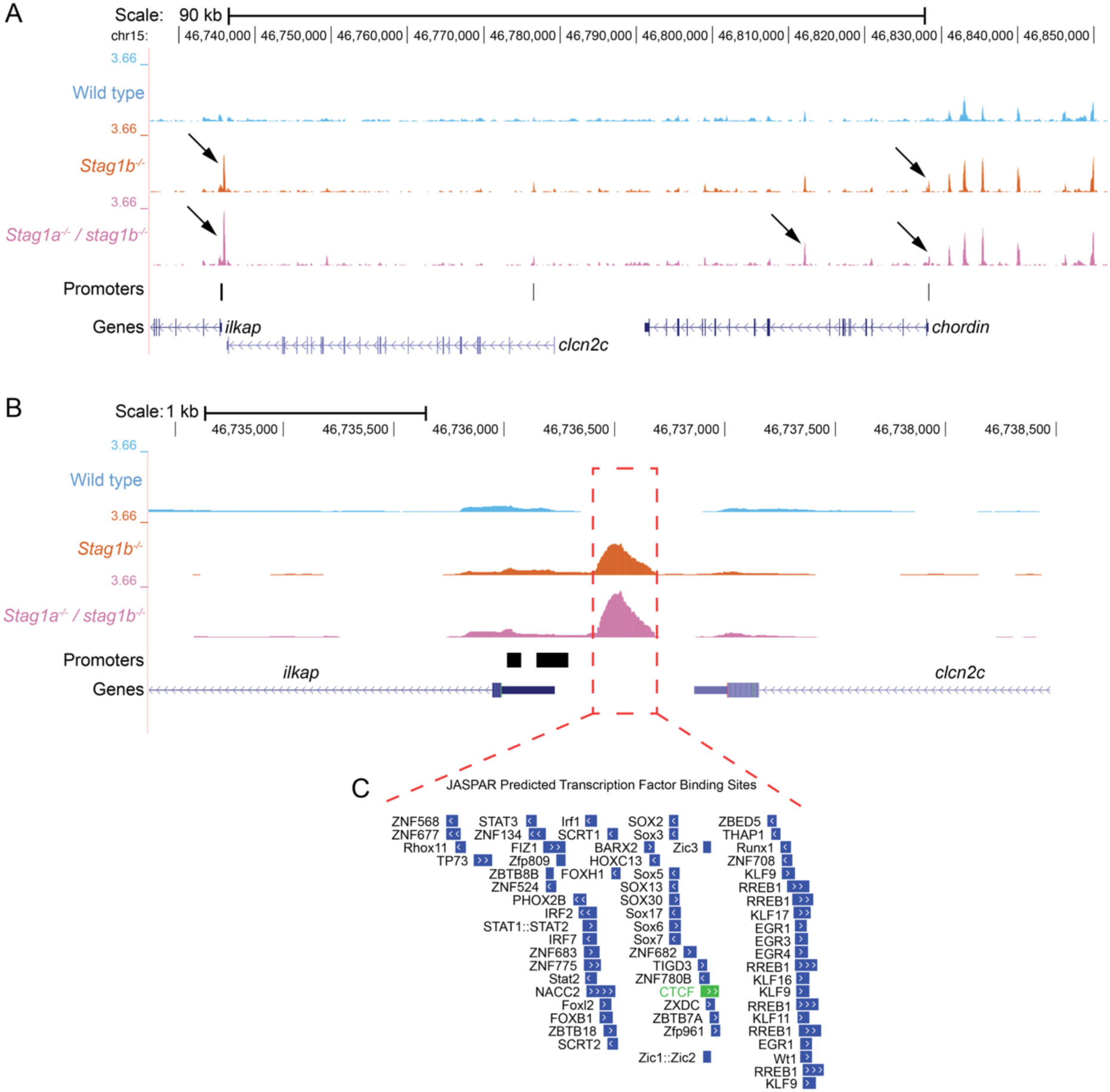
Stag1 dose-dependent increase in chromatin accessibility at an intergenic region ∼90 kb downstream of chordin in stag1 mutant tailbuds. (**A**) UCSC Genome Browser view of ATAC-seq tracks (wild type, *stag1b^-/-^,* and *stag1a^-/-^;stag1b^-/-^*) across the *chrd* locus on chromosome 15. Tracks show normalized ATAC-seq signal from 16 hpf tailbud dissections. Note the prominent *stag1* dose-dependent gain in accessibility at the intergenic region between *clcn2c* and *ilkap* (arrows left side), located approximately 90 kb downstream of the *chrd* transcription start site. Small changes in accessibility can be seen near the promoter and at intronic region of *chrd* (arrows right side). (**B**) Magnified view of the red dashed box in (A), highlighting the *stag1* dose-dependent increase in open chromatin specifically in the intergenic region. (**C**) JASPAR-predicted transcription factor binding sites within the gained accessibility peak. The region is highly enriched for motifs of developmental transcription factors (Zic, POU, KLF, SOX and homeodomain families). The CTCF motif is highlighted in green.

The region contains a CTCF motif together with binding sites for multiple developmental transcription factors (Fig. 5C). BMP signalling has been shown to induce p21 expression and promote G1 arrest in certain contexts. Therefore increased expression of the BMP antagonist *chrd* may serve to attenuate this pathway and counteract the proliferation defects caused by Stag1 loss. This model is consistent with the dose-dependent nature of both the cell cycle and *chrd* phenotypes.

To understand the mechanistic basis of these phenotypes, we next examined how loss of Stag1 affects chromatin accessibility on a genome-wide scale.

### *Stag1* loss leads to decreased chromatin accessibility near TAD boundaries and increased accessibility associated with developmental transcription factors

We identified genome-wide 886 differentially accessible regions (DARs) with gained accessibility and 576 regions with decreased accessibility in *stag1b^-/-^* mutants (5% FDR, Figure 6A, Data S4). Regions that lost accessibility were strongly enriched near TAD boundaries compared to randomly selected regions or regions that gained accessibility (Figure 6B). Motif analysis of gained accessible regions revealed strong enrichment for developmental transcription factors, particularly members of the KLF/SP family (Figure 6C). A very similar enrichment of KLF/SP motifs in gained accessible regions was observed upon CTCF knockout in zebrafish (Franke et al., 2021). We interpret this increase in accessibility at KLF/SP sites as likely indirect, resulting from weakened TAD insulation that permits developmental transcription factors to access regulatory elements that are normally protected. In contrast, decreased accessible regions were highly enriched for CTCF/BORIS motifs (18.7% of sequences, >3-fold enrichment; (Figure 6D), consistent with a role for Stag1-cohesin in maintaining accessibility at CTCF-anchored TAD boundaries.

**Figure 6.**
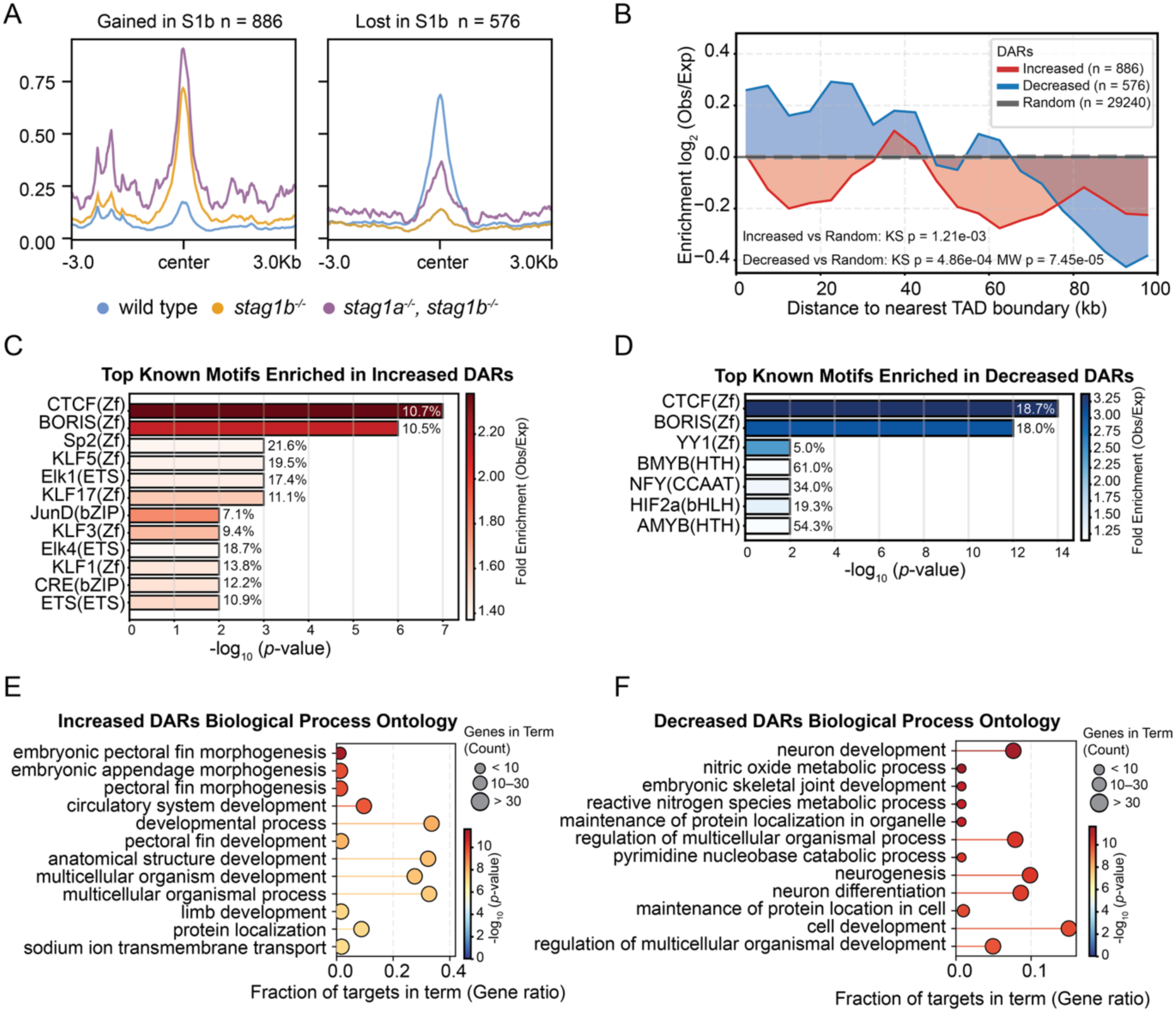
Stag1 loss leads to a decrease of chromatin accessibility at regions near TADs and increased accessibility associated with developmental transcription factors. (**A**) ATAC-seq profile of significant differentially accessible (DA) regions (5% FDR) comparing wild-type and *stag1b^-/-^* mutant tailbuds showing increased (left, n = 886) and reduced (right, n = 576) chromatin accessibility across ATAC-seq profiles form wild-type, *stag1b^-/-^* and *stag1a;stag1b* double mutants. (**B**) Log2 fold enrichment of increased and decreased differentially accessible regions in *stag1b^-/-^* tailbuds compared to random regions as a function of distance to the nearest TAD boundary. (**C**) Top known motifs enriched in DARs with increased accessibility in *stag1b^-/-^* mutants. (**D**) Top known motifs enriched in DARs with decreased accessibility in *stag1b^-/-^*mutants. Top GO terms for biological process in increased (**E**) and decreased (**F**) DARs.

We also found enrichment for B-Myb motifs among the decreased DARs which is consistent with the transcriptional repression of cell cycle progression genes (Fig. 2E) and the reduced mitotic index (Fig. 2B,C). We note that expression of MYB transcription factors themselves was largely unchanged as quantified by RNA-seq. YY1, in contrast is a direct regulator of enhancer promoter looping (Weintraub et al., 2017) that was shown to depend on cohesin to find and bind its cognate binding sites on chromatin (Hsieh et al., 2022).

To test whether the enrichment of CTCF motifs in regions of decreased accessibility reflected actual CTCF occupancy, we intersected our DARs with published CTCF ChIP-seq peaks from 24 hpf wild-type embryos (Franke et al., 2021). Decreased accessibility regions were significantly more likely to overlap a CTCF binding site than increased accessibility regions (27.1 % versus 18.6 %; odds ratio = 1.62, Fisher’s exact test P = 1.72^10̅⁴; Supplementary Fig. 4A). Nevertheless, the majority of regions that lost accessibility (73%) did not overlap a detectable CTCF peak. Consistent with a preferential effect at distal regulatory elements, both increased and decreased DARs were enriched for intergenic sequence relative to the full set of ATAC-seq peaks (75.6% and 77.6% versus 66.2%, respectively) and correspondingly depleted for intronic and promoter-proximal peaks (Supplementary Fig. 4B). Together these data indicate that Stag1 regulates chromatin accessibility through both CTCF-dependent and CTCF-independent mechanisms, acting predominantly at distal intergenic sites.

To determine the functional consequences of the altered chromatin accessibility landscapes we performed gene ontology (Biological process) enrichment analyses on genes associated with increased and decreased DARs. We found that genes linked to increased DARs were enriched for processes like embryonic morphogenesis and multicellular organism development (Figure 6E, Data S5). Decreased DARs were enriched for neurogenesis and differentiation related biological processes, as well pyrimidine catabolic processes (Figure 6F, Data S6).

### Stag1b mutation causes metabolic changes consistent with reduced mTORC1 signalling

KEGG pathway enrichment analyses showed that genes linked to increased DARs were enriched for developmental processes like embryonic morphogenesis, Notch and BMP signalling (Fig. 7A). In contrast, genes associated with lost accessibility were strongly enriched for metabolic and biosynthetic pathways, most notably mTOR signalling, aminoacyl-tRNA biosynthesis, ribosome biogenesis, and related anabolic processes (Fig. 7B). RNA-seq analysis showed a trend toward downregulation of several mTOR pathway components and regulators (*mtor, rptor, deptor, aktip*), upregulation of lysosomal genes (*atp6v0a2a* and *lamp1b*), and down-regulation of *tgfb1a*, *tgfb2* (Figure 7C, Data S7). Untargeted ¹H-NMR metabolomics of dissected tailbud tissue revealed clear changes in *stag1b^-/-^* mutants, with marked accumulation of glycerol, glycine and several free amino acids (Fig. 7C, E, Data S8). These changes, together with the downregulation of mTOR- and TGF-β-related genes indicate a shift to a more catabolic state and reduced growth in *stag1b^-/-^* mutants, consistent with the observed decrease in mitotic index (Fig. 2B-B” and C).

**Figure 7.**
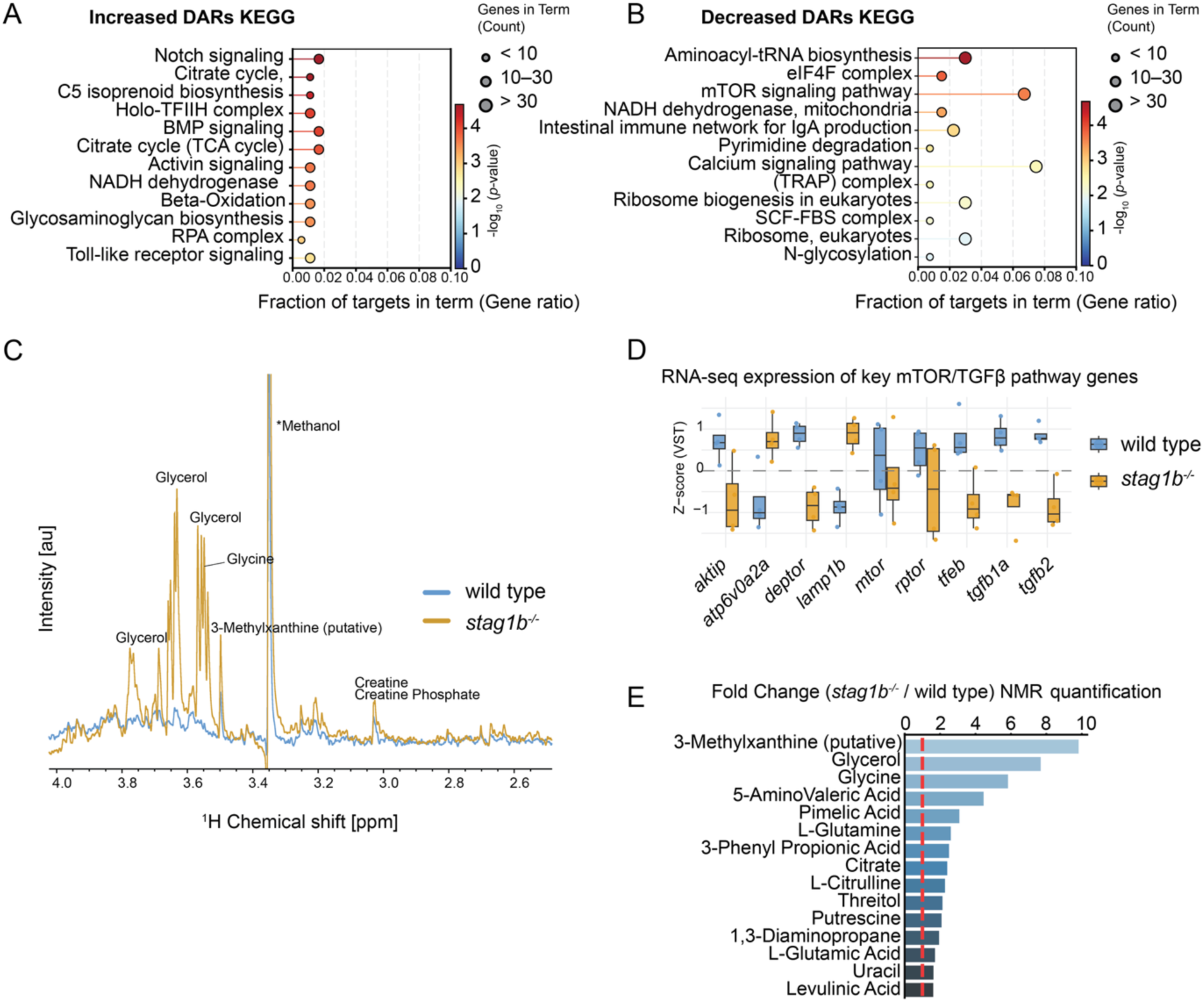
*Stag1b* mutation induces metabolic changes consistent with reduced mTOR signalling. (**A**-**B**) KEGG pathway analysis of ATAC-seq data comparing *stag1b^-/-^* with wild-type. (**A**) Increased DARs are enriched for developmental signalling pathways such as Notch and BMP while mTOR and ribosome biogenesis pathways are enriched in decreased DARs (**B**). (**C**) Overlap of two representative ^1^H-NMR spectra of tail tissue metabolites (wild-type in blue, *stag1b^-/-^*in yellow).*Methanol was excluded from the analysis as it was part of the sample processing. (**D**) RNA-seq expression of key mTOR, lysosomal and TGF-β pathway genes. (**E**) Fold-change (*stag1b^-/-^* vs. wild-type) of the most strongly altered metabolites quantified by NMR. Glycerol, glycine, and several amino acids are markedly elevated in *stag1b^-/-^* mutants.

## Discussion

Our work demonstrates that Stag1 and Stag2 perform distinct, non-redundant functions during zebrafish tailbud development. While single *stag1b* and *stag2b* mutants are viable and fertile, they exhibit markedly different transcriptional and cellular phenotypes. Consistent with our previous findings, loss of Stag2 primarily compromises Wnt signalling and mesoderm patterning with largely normal cell cycle progression. In contrast, loss of Stag1 causes broader transcriptional dysregulation, impairs cell cycle progression in a dose-dependent manner, and activates the p53 pathway. These cell cycle defects occur without the catastrophic mitotic failure seen in *rad21* or double homozygous *stag1b;stag2b* mutants. Stag1 has been implicated in telomere replication (Remeseiro et al., 2012). More recent work has shown that Stag1 plays a major role in sister telomere cohesion that is largely independent of the core cohesin ring and distinct from Stag2 (Lin et al., 2016). In addition, STAG proteins including Stag1 have been shown to interact with RNA and R-loops and to promote cohesin loading even in the absence of the cohesin ring (Porter et al., 2023). Together, these findings suggest that Stag1 contributes to genomic stability in ways that extend beyond its canonical role in sister chromatid cohesion.

A particularly striking difference between the two subunits lies in their effects on developmental signalling. Unlike *stag2b* mutants, *stag1* mutants show specific dysregulation of the BMP pathway, including dose-dependent upregulation and anterior expansion of the BMP antagonist *chrd* together with reduced *bmp4* expression. One possible interpretation is that increased *chrd* expression represents a compensatory response aimed at attenuating BMP signalling, which can promote p21 expression and cell cycle arrest in certain contexts. This could partially counteract the proliferation defects caused by Stag1 loss. This model is consistent with the dose-dependent nature of both the cell cycle and *chrd* phenotypes, although future genetic experiments will be required to test this relationship directly.

To investigate the mechanistic basis of these phenotypes, we examined genome-wide changes in chromatin accessibility. Regions that lost accessibility in *stag1* mutants were strongly enriched near TAD boundaries and for CTCF motifs, consistent with a preferential role for Stag1 at TAD boundaries (Cuadrado et al., 2019). In contrast, regions that gained accessibility were enriched for motifs of developmental transcription factors, particularly members of the KLF/SP family, a pattern also observed upon CTCF knockout in zebrafish (Franke et al., 2021). We interpret the increased accessibility at KLF/SP sites as likely indirect, resulting from weakened TAD insulation. Decreased DARs overlapped experimentally determined CTCF binding sites more frequently than increased DARs; nevertheless, the majority of accessibility losses occurred outside detectable CTCF peaks. Together these observations indicate that Stag1 influences chromatin accessibility through both CTCF-associated and CTCF-independent mechanisms.

Locally, at the *chrd* locus, we observed smaller increases in accessibility near the gene itself alongside a much stronger *stag1* dose-dependent gain at a distal intergenic region ∼90 kb downstream. The region contains a CTCF motif together with binding sites for multiple developmental transcription factors.

Together, these results indicate that Stag1 loss triggers a combination of defects in chromatin architecture, cell cycle progression and developmental signalling. The upregulation of *tp53* and p21, together with a shift toward a more catabolic metabolic state, likely reflects cellular responses to impaired TAD insulation and replication stress. These findings establish that Stag1 and Stag2 are not merely interchangeable cohesin subunits but perform distinct, partially independent functions in chromatin architecture, cell-cycle control, and developmental gene regulation. The pronounced differences between single *stag* mutants and core cohesin (*rad21*) mutants highlight the importance of studying individual cohesin accessory subunits to understand both normal development and the tissue-specific consequences of cohesin mutations in human disease.

## Methods

### Zebrafish Husbandry

Wild type (WIK) (Rauch et al., 1997), stag1anz204 (Ketharnathan et al., 2020), stag1bnz205 (Ketharnathan et al., 2020), stag2bnz207 (Ketharnathan et al., 2020) and rad21nz171 (Horsfield et al., 2007) zebrafish lines were maintained at 28 °C according to established husbandry methods (Westerfield, 1995). Zebrafish were housed in the Otago Zebrafish Facility (Department of Pathology, University of Otago, Dunedin, New Zealand). All animal work was performed in accordance with the Otago Zebrafish Facility Standard Operating Procedures (AUP 21-110) and under Environmental Risk Management Authority approval numbers GMC005627, GMD100922 and GMC001366. For all experiments, embryos were developed at 20 or 28 °C.

### Tailbud bulk RNA sequencing (RNA-seq) and analyses

Tailbuds were dissected from stage-matched embryos at 16 somites (16-18 hpf) as illustrated in Fig. 1A. For RNA-seq, tailbuds were individually lysed in 3 μL of RLT + BME (Qiagen RNeasy) and stored in separate PCR tubes at -80 °C. Total RNA was extracted from the pools of 80 tailbuds per sample using the RNeasy Micro kit (74104; Qiagen, Germany). Quality and quantity of RNA were assessed using Qubit 4.0 Fluorometer (Thermo Fisher Scientific, USA), Agilent RNA 6000 Nano Kit on 2100 Bioanalyzer (Agilent Technologies, Netherlands) and NanoPhotometer NP80 Touch (Implen GmbH, Germany).

Libraries were prepared from 250 ng of total RNA using the TruSeq Stranded mRNA Library Prep kit (Illumina, USA) and TruSeq RNA CD Index Plate (Illumina, USA) for sample multiplexing. The concentration of the libraries was quantified using a Qubit 4.0 Fluorometer (Thermo Fisher Scientific, USA), and the mean fragment size was assessed using the DNA High Sensitivity KIT on a 2100 Bioanalyzer (Agilent Technologies, Netherlands). 4 nM pooled libraries was sequenced on NovaSeq S1 flow cell through Genohub.

RNA-seq reads were trimmed using Cutadapt (Martin, 2011), and aligned to the reference genome (GRCz11.113) and a curated GTF file obtained from (Lawson et al., 2020) with STAR (Dobin et al., 2013). FeatureCounts (Liao et al., 2014) was used to generate fragment count matrices. DESeq2 (Love et al., 2014) was used to perform differential gene expression analysis, and multi-testing correction was done using the Benjamini-Hochberg procedure. The false discovery rate (FDR) threshold was set at 5%. Enrichment analysis were performed using clusterProfiler (Yu et al., 2012) against the background of all expressed genes above a minimum threshold.

### Immunohistochemistry - pHH3 staining

To detect cells in mitosis, fixed embryos (4% PFA at 4 °C overnight) were stained for phosphorylated histone H3 (pHH3). Embryos were rehydrated through a graded methanol/PBST series, washed twice in 100% PBST, then washed three times in sterile deionised water. Embryos were permeabilised in acetone at –20 °C for 10 min, washed again in deionised water, and blocked in blocking solution (2% FBS in PBS).

Embryos were incubated overnight at 4 °C with rocking in anti-pHH3 rabbit primary antibody (#3377, Cell Signalling Technology) diluted 1:1000 in blocking solution. The following day, embryos were washed three times for 10 min in PBST, followed by two 10-min washes in 1% fetal bovine serum (FBS) in PBST. Embryos were then incubated for 2 days at room temperature (protected from light) with goat anti-rabbit Alexa Fluor 488 secondary antibody (#A11008, Thermo Fisher Scientific) diluted 1:1000 in 1% FBS/PBST.

After incubation, embryos were washed five times for 10 min in PBST (protected from light) and imaged using a Nikon C2 confocal microscope as z-stacks of the optical sections. Maximum-intensity projections were generated from 33 optical sections. The look-up table was set to 300 for pHH3 and kept constant across all samples.

### Quantification of pHH3-positive cells

pHH3-positive cells were quantified in ImageJ. Images were converted to 8-bit, inverted, and thresholded at a darkness value of 140. A 200 × 200-pixel square was placed over the posterior tailbud region, and positive cells within this area were counted using the “Analyze Particles” function. The number of pHH3-positive cells per tailbud was recorded for each genotype.

### Flow cytometry

Embryos at the 16-somite stage were fixed in methanol and tailbuds (n=30) were dissected. Cells were rehydrated in PBS, filtered through a 40 μm cell strainer and nuclei were stained with DRAQ5 (#ab108410, Abcam) at 5 μM final concentration on ice for 45 min in the dark. Cell cycle profiles of three independent replicates for each genotype were obtained using a BD FACS Aria III (BD Biosciences). Data analysis and plots were generated using Cytoflow (Teague, 2022).

### Western blot

Around 30 embryos staged at 16 somites were lysed in RIPA buffer supplemented with protease and phosphatase inhibitors (cOmplete, Merck). Equal amounts of protein (20 μg) were separated by SDS-PAGE, transferred to nitrocellulose membranes, and blocked in EveryBlot blocking buffer (Bio Rad). Membranes were incubated overnight with anti-Cdkn1a (1:1000) (#2947, Cell Signalling Technology) and anti-α-Tubulin (1:5000) (ab7291, Abcam) antibodies, followed by fluorescently labelled secondary antibodies (1:5000) (StarBright Blue, Bio Rad). Wild-type embryos exposed to UV irradiation served as a positive control for p53 pathway activation.

### Quantitative RT-PCR

Complementary DNA (cDNA) was synthesised from 1 µg of total RNA (dissected tailbuds) using the qScript cDNA SuperMix (Quantabio) according to the manufacturer’s instructions. RT-PCR was performed in 10 µL reactions containing 1 µL diluted cDNA, gene-specific primers (see Data S1) and PowerUp SYBR Green Master Mix (Thermo Fisher Scientific) on a QuantStudio 5 Real-Time PCR System (Thermo Fisher Scientific). Cycling conditions were: 2 min at 50 °C, 2 min at 95 °C, followed by 40 cycles of 15 s at 95 °C and 1 min at 60 °C. A melt curve was generated after each run to confirm specificity. Relative expression was calculated using the ΔΔCt method with *eef1a1l1* and *rpl13a* as the internal controls. Data were analysed with QuantStudio Design & Analysis Software and statistical significance was determined by one-way ANOVA with appropriate post-hoc testing.

### Whole-mount in situ hybridisations (WISH)

WISH for *chrd and bmp4* was performed using 0.5 ng/μL of riboprobe as previously described (Kalev-Zylinska et al., 2002). Primers used to clone cDNA fragments are listed in Data S9.

### ATAC-seq cell isolation and tagmentation

ATAC assay was performed in duplicates on 50,000 cells per technical replicate as described previously in the Omni-ATAC protocol (Corces et al., 2017) with some modifications. Tailbuds were dissected in PBS and cells were spun down at 500g at 4 °C and resuspended in ATAC-seq resuspension buffer (RSB) (10 mM Tris-HCl pH 7.4, 10 mM NaCl, and 3 mM MgCl_2_ in water). Cells were spun again for 5 min at 500g and resuspended in ATAC-seq RSB containing 0.1% NP40, 0.1% Tween-20, and 0.01% digitonin (Cell Signalling, #16359) by pipetting up and down three times. The cell lysis reaction was incubated on ice for 3 min. After lysis, 1 ml of ATAC-seq RSB containing 0.1% Tween-20 (without NP40 or digitonin) was added, and the tubes were inverted to mix. Nuclei were then centrifuged for 10 min at 500 *g* at 4 °C. Supernatant was removed by carefully avoiding the nuclei pellet. Nuclei were suspended in 50 μl of transposition mix (25 μl of 2× TD buffer (20 mM Tris-HCl pH 7.6, 10 mM MgCl_2,_ 20% v/v Dimethyl Formamide), 2.5 μl assembled Tn5 (Cat#. 53150, Active Motif), 16.5 μl PBS, 0.5 μl 1% digitonin, 0.5 μl 10% Tween-20, and 5 μl water) by pipetting up and down six times. Transposition reactions were incubated at 37 °C for 30 min in a thermomixer with shaking at 800 r.p.m. Reactions were cleaned up with Monarch PCR & DNA cleanup columns. PCR amplification of libraries using standard Illumina/Nextera i5 and i7 compatible indexing primers was done as previously described (Buenrostro et al., 2015). Care was taken not to overamplify libraries by running an aliquot RT-PCR and stopping after 1/3 of maximum fluorescence was reached, usually around 12 total cycles. Prior to sequencing, fragment length distribution and DNA concentration were determined via PAGE and Qubit respectively. Libraries were pooled to 4 nM and sequenced on a NovaSeqX to at least 100 million paired end reads (150 bp) per sample.

### ATAC-seq analysis

Raw reads were processed using the nf-core/atacseq pipeline (v2.1.2) (Patel et al., 2023) which handles quality control, alignment, normalisation, peak calling, and quantification. After quality and adapter trimming reads were mapped to the zebrafish genome (GRCz11). For differential peak accessibility analysis we used a sliding window approach with csaw (Lun & Smyth, 2016) and DESeq2 (Love et al., 2014) with a 5% FDR cutoff. Normalised bigwig files of aligned reads were visualised with UCSC genome browser (Casper et al., 2026).

### TAD boundary proximity and enrichment analysis of differentially accessible regions

TAD boundaries were taken from publicly available Hi-C data (Wike et al., 2021) through the 4D Nucleome Data Portal (Reiff et al., 2022). Distances from the centre of each DAR (and from size- and chromosome-matched random intervals) to the nearest TAD boundary were calculated with bedtools closest (Quinlan & Hall, 2010).

Two statistical tests were applied to the resulting distance distributions: two-sample Kolmogorov–Smirnov tests (to compare the full empirical distributions) and Mann–Whitney U tests (to compare central tendency). Both tests were performed in Python using the SciPy (Virtanen et al., 2020) stats packages scipy.stats.ks_2samp and scipy.stats.mannwhitneyu. For visualisation of distance-dependent enrichment distances were binned into 5 kb intervals from 0 to 100 kb. Bin fractions were computed for gained (frac_inc), decreased (frac_dec) and random (frac_rand) sets. A pseudocount of 1 × 10̅⁶ was added to every fraction, and log₂ enrichment was calculated as:

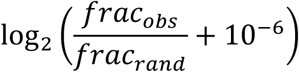

The resulting enrichment profiles were smoothed with scipy.ndimage.uniform_filter1d (window size = 5 bins, mode = ‘nearest’) (Virtanen et al., 2020) and plotted against distance to the nearest TAD boundary.

Motif enrichment was performed with HOMER (Heinz et al., 2010) on sequences centred on gained and decreased DARs (±250 bp) using GC-matched random genomic regions as background. Gene Ontology (Biological Process) and KEGG pathway over-representation analysis was performed with clusterProfiler (Yu et al., 2012) on genes associated with the DARs; terms with FDR < 0.05.

### NMR spectroscopy analysis

Approximately 30 tailbuds from 16 hpf embryos were prepared and stored at -80 °C until processing. Samples were thawed on ice and homogenized by repeated aspiration through a 25-gauge needle. Ice-cold 98% methanol (1 mL) was added to promote metabolite extraction and protein precipitation, and samples were incubated on dry ice for 20 min. Following centrifugation, the supernatant was collected and dried under vacuum for 2.5 hours. Dried extracts were stored at −80 °C until analysis. Prior to measurements, samples were reconstituted in 540 µL phosphate buffer (50 mM, pH 7.4) and 60 µL D2O and transferred to a 5 mm Bruker SampleJet NMR tube. One-dimensional ^1^H NMR spectra were obtained using a Bruker 600 Mhz spectrometer operated by an Avance III console equipped with a TXI triple-resonance probe including z-gradients. A WATERGATE pulse sequence with W5 pulse elements was applied by acquiring 1024 scans, an acquisition time of 2 s, and a relaxation delay of 2 s at 298 K. A total of 16384 complex data points were acquired and a 2 Hz exponential apodization function was applied during processing. Upon referencing of the spectra, baseline and phase corrections were applied using TopSpin NMR Suite (Bruker BioSpin GmbH, v4.5.0). Metabolite identification and quantification were carried out with Chenomx NMR Suite (Chenomx Inc., v9.0), which allows the qualitative and quantitative analysis of NMR spectra by matching spectral features to a reference database.

## Data availability

The data that support this study are available from the corresponding author upon reasonable request. The RNA-seq and ATAC-seq data generated in this study have been deposited in the Gene Expression Omnibus (GEO) database under accession code GSE342623. The public datasets used in this study are available in the GEO database under accession codes: GSE152744 and GSE156096.

## Acknowledgements

The authors would like to thank Dr Doug Mackie for expert management of the zebrafish facility.

## Supplementary Figures

**Supplementary Figure 1.**
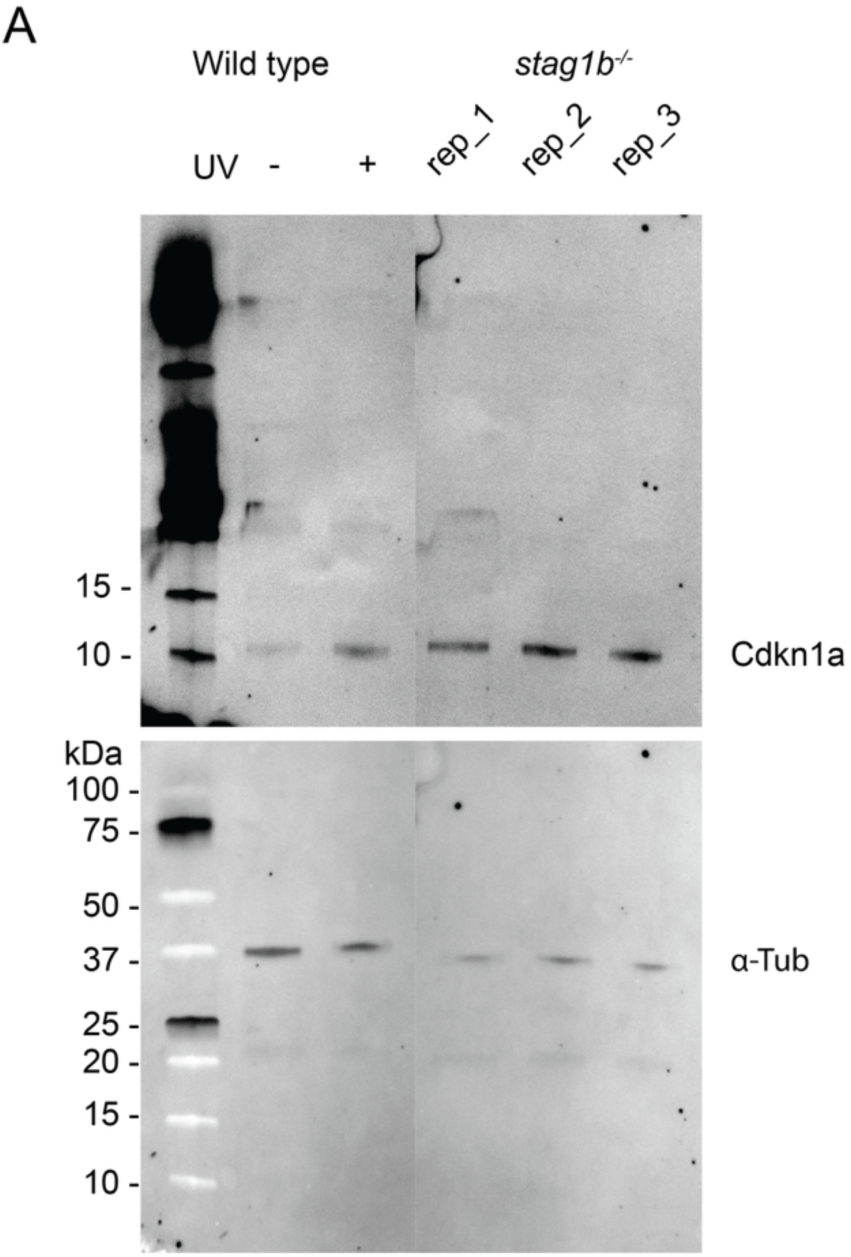
Western blot analysis of Cdkn1a (p21) protein levels in wild-type and *stag1b^-/-^* tailbuds. (**A**) Representative Western blot for Cdkn1a (upper panel) and α-Tubulin loading control (lower panel). Wild-type embryos were either untreated or treated with UV (positive control for DNA damage-induced p53 activation). *Stag1b^-/-^*samples are three independent biological replicates of untreated tailbuds at 16 hpf. Molecular weight markers (kDa) are shown on the left.

**Supplementary Figure 2.**
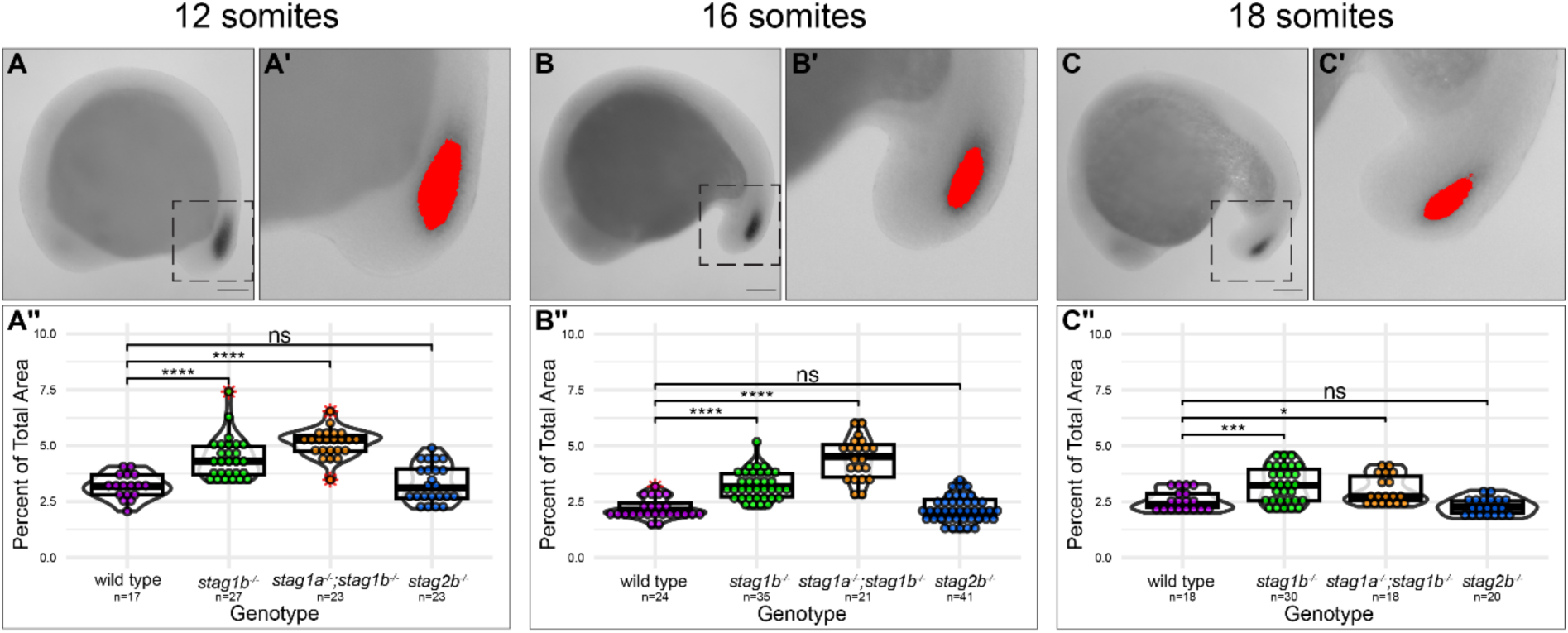
*Chrd* expansion in *stag1^-/-^* mutants is not due to developmental delay. Tailbud *chrd* area was measured in *stag* mutant and wild-type tailbuds at 12 somites (**A-A’’**), 16 somites (**B-B’’**) and 18 somites (**C-C’’**). The boxes in A, B and C represent the cropped images in A’, B’ and C’ respectively, where the area of *chrd* staining was measured. Violin and box plots show the percentage of *chrd* positive pixels in wild-type and *stag* mutants at 12 somites (**A’’’**), 16 somites (**B’’’**) and 18 somites (**C’’’**). Statistical significance was determined using a t-test comparing the mean tailbud area of *chrd* in wild type and each mutant individually: *p* * ≤ 0.05, *p* *** ≤ 0.001, *p* **** ≤ 0.0001, ns = not significant.

**Supplementary Figure 3.**
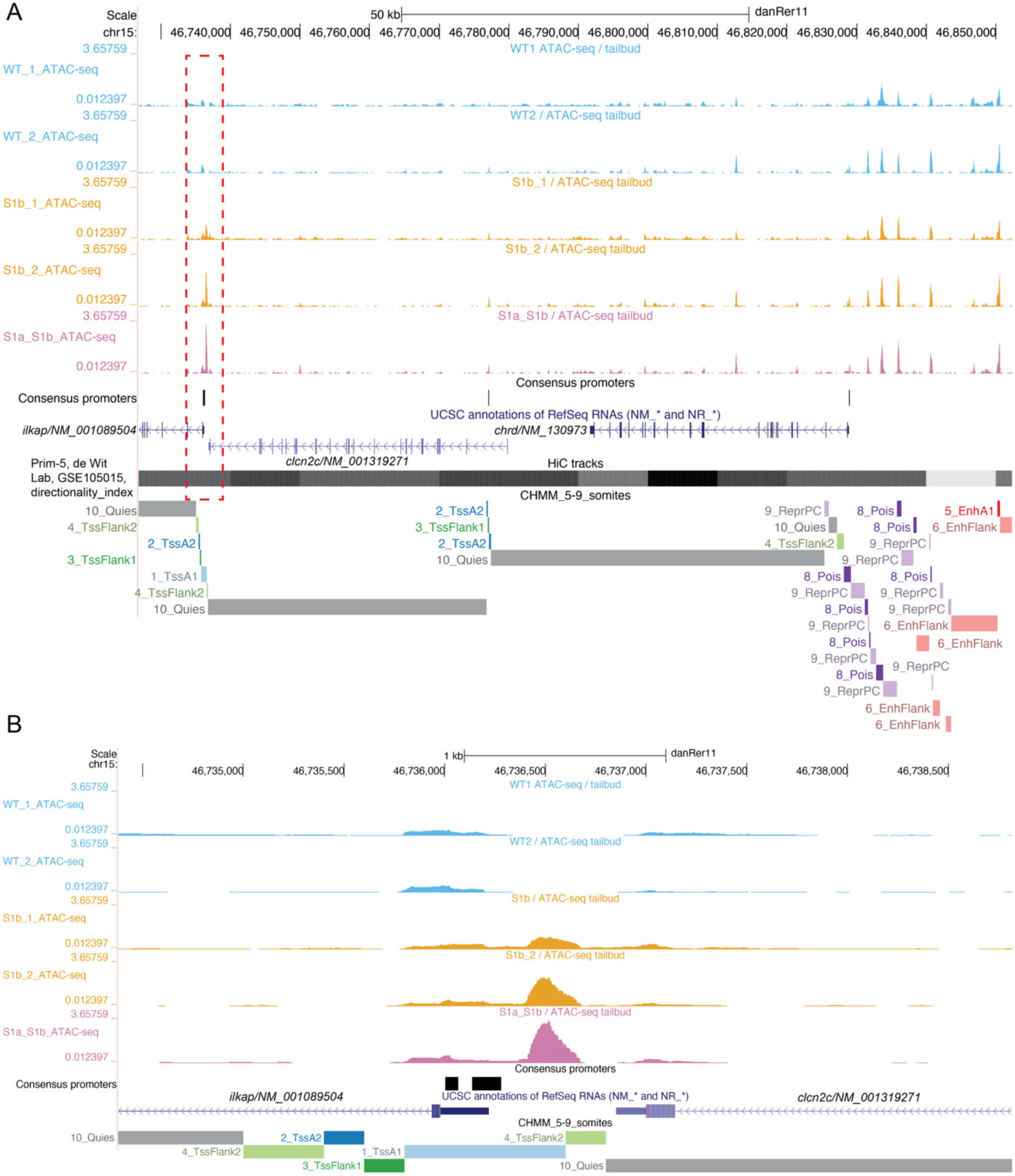
Stag1 dose-dependent increase in chromatin accessibility at a distal intergenic region downstream of *chrd*. (**A**) UCSC browser view of wild-type *stag1b^-/-^* and *stag1a^-/-^/stag1b^-/-^* ATAC-seq data and chromatin states at the *chrd* locus. (**B**) Magnified view of red dotted area in (**A**) showing a stag1 dose dependent increase in open chromatin at the intergenic region between *clcn2c* and *ilkap* ∼90 kbp downstream of *chrd*.

**Supplementary Figure 4.**
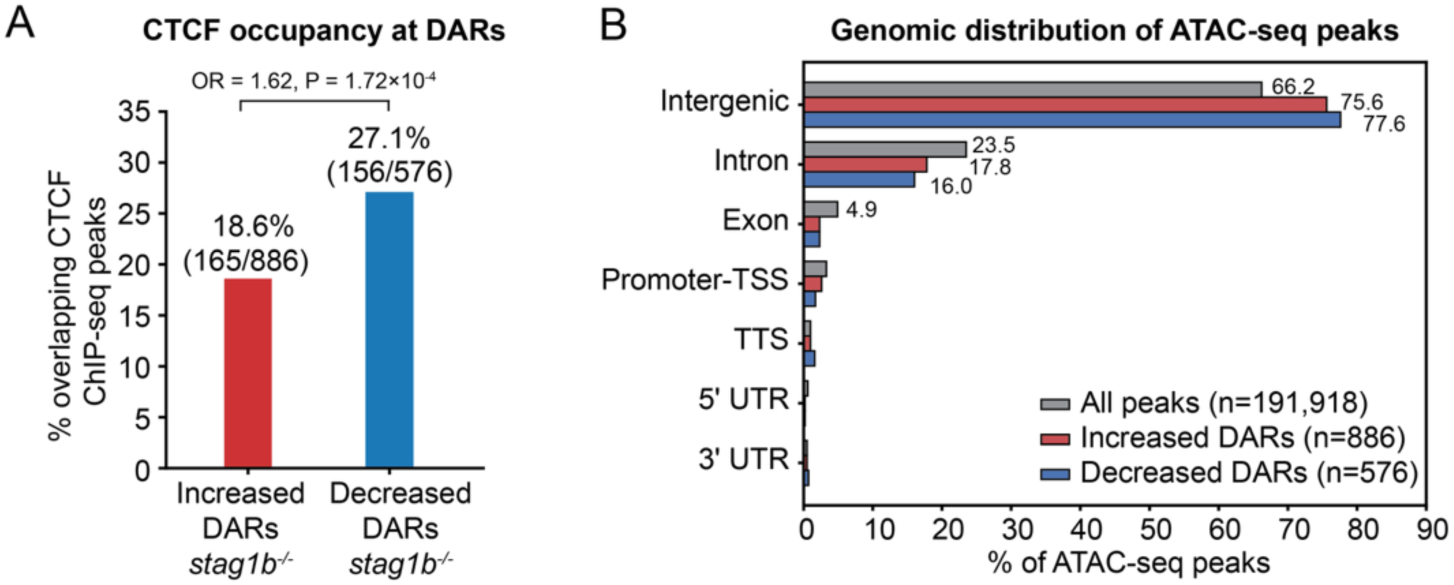
CTCF occupancy and genomic distribution of differentially accessible regions in stag1b mutants. (**A**) Percentage of increased and decreased DARs that overlap CTCF ChIP-seq peaks from 24 hpf wild-type embryos (Franke et al., 2021). Decreased DARs are significantly more likely to overlap a CTCF binding site than increased DARs (27.1 % vs 18.6 %; Fisher’s exact test, odds ratio = 1.62, P = 1.72^10⁻⁴). (**B**) Genomic distribution of all consensus ATAC-seq peaks, increased DARs and decreased DARs. Both classes of DARs are enriched for intergenic sequence relative to the overall peak set.

